# Nematic cell alignment directs calcium waves

**DOI:** 10.1101/2025.10.20.683494

**Authors:** Annemarie C. Winterstrain, Bennett C. Sessa, Michael M. Norton, Hannah G. Yevick

**Affiliations:** Department of Physics, Brandeis University, Waltham, Massachusetts 02453, USA

## Abstract

Tissues rely on supracellular signals to coordinate their cells over a long range. Two such tissue-scale cues are calcium waves and patterns of cell-cell alignment or nematic order. During wound healing, for example, calcium waves propagate across a tissue to guide directed cell migration and reepithelialization. Defects in long-range cell-cell alignment, or nematic orientation, can act to localize morphogenetic events in a tissue. Although these two cues have been considered in isolation, we demonstrate a relationship in epithelial tissue between long-range calcium signaling and the cell’s nematic order: The speed of a wound-induced calcium wave depends monotonically on the angle between the wave vector and cell axis, with maximal wave speed occurring perpendicular to the tissue’s orientation. Including anisotropic diffusive coupling between cells in a canonical reaction-diffusion model recapitulates our measured calcium wave dynamics. Our model demonstrates how orientation defects can desynchronize information propagation across a tissue. A calcium wave front is bent around nematic defects, therefore cells the same distance from a wound can receive the calcium signal at different times. Our work elucidates how spatial patterns in global cell alignment can control collective communication via calcium signaling during development, wound healing, and disease.

---

Cells rely on supracellular cues to collectively coordinate tissue-scale behaviors. Tissue-scale calcium waves are one such signal. Intercellular calcium waves appear during the synchronized movements during wound healing [1, 2] and tissue development [3–6]. Alternatively, supracellular cues can be mechanical, where the tissue’s local mechanics can relay information to the cell about its environment. For example, anisotropic stresses caused by a tissue’s nematic structure [7] can position within a tissue important biological phenomena including apoptosis [8], limb budding [9–11], and accumulation of neural progenitor cell defects [12]. Many of these phenomena that are co-local with defects in a tissue’s nematic order, including cell death, cell extrusion and limb development, simultaneously exhibit calcium spikes [13–15]. Yet, how calcium signaling and nematic alignment establish collective cues in epithelia has only been considered in isolation.

A calcium flux can be triggered in a cell from a mechanical stimulus (e.g., wounding) or a chemical signal [1, 16]. When a sufficiently large increase in intracellular Ca^2+^ is triggered, an intercellular Ca^2+^ wave can propagate. Depending on the biological context, global wave moves through tissue in two ways: intercellularly through gap junctions or through released extracellular signals such as ATP [17].

Local cell-cell coupling and tissue-scale structure are inherently connected to cell shape and cell-cell alignment. At the tissue scale, the degree of cell-cell alignment can vary. Elongated cells can organize like liquid crystals, forming locally well-aligned ’nematic’ domains [7, 18]. The active nature of epithelia spontaneously creates disorder in the form of motile topological defects, which has been well-captured through the framework of active nematohydrodynamics [7, 8, 19, 20]. Anisotropy is not only important for mechanics, but also for signal propagation.

For example, it is well appreciated that highly specialized cells, such as myocytes, leverage their anisotropy to directionally propagate signals at the tissue scale and perform their functions [21]. Yet, how nematic order couples to the propagation of calcium waves in epithelial tissues is unknown.

We show that the speed of calcium waves across an epithelial tissue depends on the angle between a cell’s local nematic alignment and the orientation of the wave. We found that elongated RPE-1 cells have increased connectivity laterally with their co-aligned neighbors, leading to calcium waves propagating fastest *perpendicular* to the tissue’s nematic director. Further, we found that chemically inhibiting gap junctions and increasing cell density, respectively, decrease and increase wave speed. We frame our experimental observations with a simple model that posits that the propagation of calcium signals from one cell to its neighbors is dominated by anisotropic cell-cell coupling. Our model is able to recapitulate our experimental results, as well as predict how calcium waves move in tissues with nematic defects. Waves emanating from point wounds on aligned tissue evolve into ellipsoidal contours, whereas point wounds on nematic defects inherit symmetries from the underlying nematic: trefoil for − 1/2 defects and circular profiles with a translating center of mass for +1/2. Our results show how a tissue’s nematic alignment, not spatial distance from a wound alone, controls when a cell receives a calcium signal. Our results demonstrate the importance of considering cellular nematics when interpreting how information propagates across epithelial tissue. Our quantitative description of how topological defects impact calcium wave movement elucidates how diverse developing tissues that exhibit calcium waves during morphological changes and wound healing collectively communicate.

## RESULTS

Retinal pigment epithelial cells (RPE-1) are an ideal cell model to investigate the interplay between nematic cell alignment and calcium wave propagation in an epithelial tissue. RPE-1 cells have a rod-like morphology making their orientation easily apparent. RPE-1 cells have a tendency to locally orient with their neighbors. Cell-cell interactions lead to neighborhoods of aligned cells but confluent unconfined RPE-1 tissue remains globally disordered (Fig. 1*A*) [7, 22]. We engineered longrange nematic order in the tissue by seeding cells on micro-grooved substrates. Cells on these substrates all oriented their long axis with the direction of the microgrooves (Fig. 1*B*) suppressing the nematic defects seen on flat smooth substrates. Aligned cells still retain a characteristic epithelial packing pattern with an average of 6 neighbors (Fig. S1*J*). With purely aligned tissues, we could address whether the relative angle between the cell’s nematic director 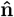 and the direction of wave propagation impacts the wave speed (Fig. 1*I*).

**FIG. 1.**
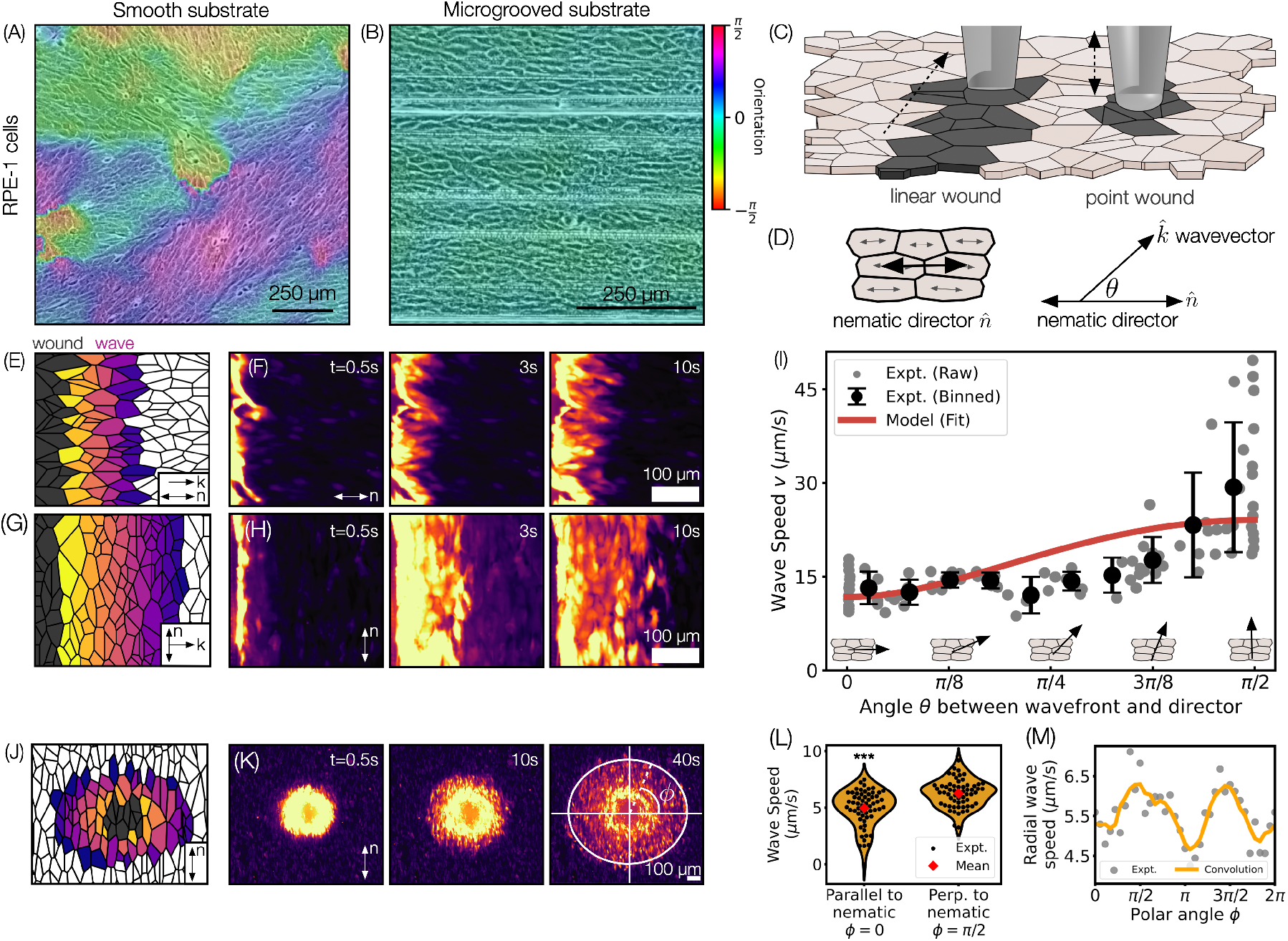
Nematic director controls speed of calcium waves initiated by linear and point wounds. (A,B) Phase contrast images of RPE-1 cells grown on a plastic culture dish (A) and a microgrooved substrate (B), overlaid with orientation fields. (C) Schematic illustrating experimental wounding methods. Linear wounds were created by dragging a pin across the tissue, while localized wounds were produced by gently tapping the surface with a rounded tool. (D) Schematic illustrating the nematic director 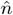, which lies along the aligned cell’s long axis, and the wavevector 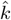 from the propagating calcium signal. We define *θ* as the angle (from 0 to 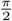) between the nematic director and the wavevector. (E) Schematic and (F) time series of a linear wound which initiated a wave traveling parallel to the nematic director (*θ* = 0). (G) Schematic and (H) time series of a linear wound which initiated a wave traveling perpendicular to the nematic director (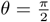). (I) Experimental measurements of wave speed (gray disks) as a function of orientation *θ*. Error bars represent the standard deviation of the binned wave speeds. Waves travel significantly faster perpendicular (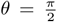) than parallel (*θ* = 0) to the nematic director. Red line represents the model. Cell diagrams demonstrate the wave propagation direction with an arrow, relative to the tissue organization. (J) Schematic and (K) time series showing outward wave propagation from a localized wound. White diagram on *t* = 40s image defines the polar angle *ϕ*. When the polar angle *ϕ* = 0, the wave at that point in the contour is moving parallel to the nematic director. When the polar angle 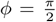, the wave at that point in the contour is moving perpendicular to the nematic. (L) Quantification of wave speed from a localized wound in the *ϕ* = 0, 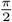 directions. Waves propagate faster in the 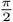 direction. *** indicates *p<* 0.001 using a paired t-test. Wave speeds have been averaged radially. (M) Radial profile of wave speed from a localized wound. The wave travels fastest perpendicular to the nematic axis 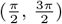.

The aligned tissues were incubated with Calbryte-520AM a cell permeable fluorescent indicator of intra-cellular calcium levels. We scratched a line across the plate of cells with a sewing pin to create linear wounds and measured the changes in fluorescence in the cells over *∼*1 min. Linear wound experiments were performed at different angles relative to the global nematic orientation 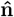 (Fig. 1*D*). The mechanical wounding of the tissue resulted in a calcium flux in cells next to the wound and subsequent outward propagation of a calcium wave front. For linear wounds, a tissue-scale linear wave moved out perpendicular to the scratch (Fig. 1*F, H*). We identified that the speed of the calcium wavefront depended on the cell’s orientation with respect to the incoming wave. The wave speed is maximal when the wound is aligned along the nematic director (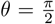, i.e., scratch parallel to the cell’s long axis), and minimal when the wound is perpendicular to the director (*θ* = 0, scratch along the cell’s short axis), with intermediate speeds for oblique orientations (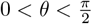) (Fig. 1*I*).

We next asked if the speed of a radial wavefront exhibited the same dependence on cell nematic orientation as a linearly propagating wavefront. We locally wounded a uniformly aligned nematic tissues with a small spherical tip to create point wounds (Fig. 1*K*). Similarly to linear wounds, point wounds triggered cells adjacent to the wound to light up and subsequently a wave propagated away from the wound. Waves traveled slower in all directions compared to the linear wound. Still, the calcium wave progressed faster perpendicular to the nematic compared to parallel to the nematic (Fig. 1*L, M*).

Next, we investigated the origin of the anisotropy in the speed of a calcium wave across an aligned tissue. We hypothesize that wave speed anisotropy arises from greater connectivity between the lateral sides of elongated cells. Schematically, we illustrate the logic behind our hypothesis in (Fig. 2*A*), showing how increased contact area naturally leads to stronger coupling through a greater number of gap junctions. We directly explored cell-cell connectivity with immunofluorescence staining of Connexin-43, a gap junction protein, confirming that cells were preferentially connected laterally by a greater quantity of junctions along the cell’s long axis compared to the cell’s short axis (Fig. 2*B*).

**FIG. 2.**
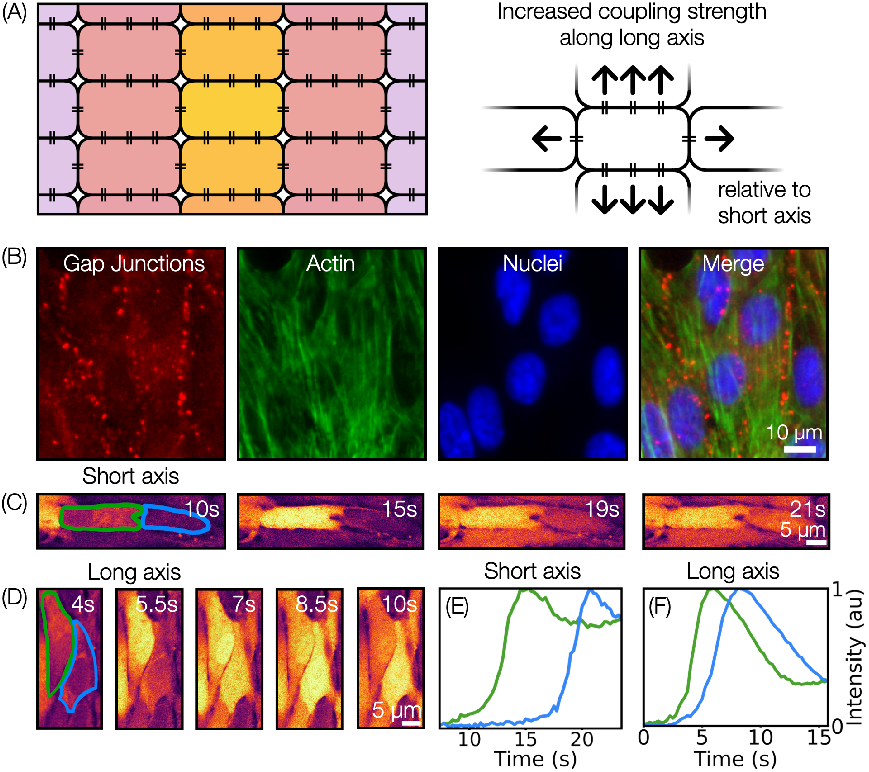
Gap junction density determines intercellular coupling and wave speed. (A) [Left] Schematic demonstrating cells (black rectangles) in a tissue with evenly dispersed gap junctions (double lines) along their perimeter. Each cell has a higher proportion of gap junctions along their long axes than their short axes. [Right] Zoomed in schematic of one cell demonstrating that the long axis has an increased coupling strength relative to the short axis given it contains more gap junctions. (B) Immunofluorescence image of a cell monolayer showing gap junctions (red), actin filaments (green), and nuclei (blue). Gap junctions are distributed more densely along the long axis compared to the short axis. (C) Time series of wave propagation across two tip-to-tip cell neighbors. The calcium waves moves across the short axis of the cell. (D) Time series of wave propagation across two lateral cell neighbors. The calcium wave moves across the long axis of the cell. (E,F) Fluorescence intensity profiles over time for panels (C) and (D), respectively. The time between intensity peaks is longer for waves moving across the short axis of the cell than the long axis.

We further quantified anisotropy at the cell scale by examining the calcium dynamics of pairs of cells that are either connected end to end (Fig. 2*C*) or connected side to side (Fig. 2*D*). We observe that in both cases, the interior of a given cell activates nearly uniformly. That is, upon stimulation, internal wave propagation is rapid – much faster than the wave speed at the tissue scale; we further quantify this in the supplement (Fig. S2). Strikingly, however, the time between spikes for the two pairs is different. End-to-end connected cells show a large time lag between spikes compared to that of the lateral pair. Fig. 2*E,F* plots the cells’ intensity at their centers over time, demonstrating this difference. From this, we conclude that gap junctions control the time lag of calcium spikes between cells, thereby determining the wave speed at the tissue scale.

To test the importance of gap-junction-mediated coupling at the tissue scale, we experimentally decreased gap-junction activity and measured wave speed. We titrated the inhibition of gap junctions using Meclofenamic Acid which is reported to block gap junction communication in RPE cells [23]. A linear wound was performed at an angle of 0 radians. At concentrations of 5*μ*M and 15*μ*M the wave can still propagate, but it moved slower than in the control where no inhibitor is added (Fig. 3*A*). At and above 25*μ*M we did not observe any tissue-scale waves and only the cells that contacted the wound lit up (Fig. S3).

**FIG. 3.**
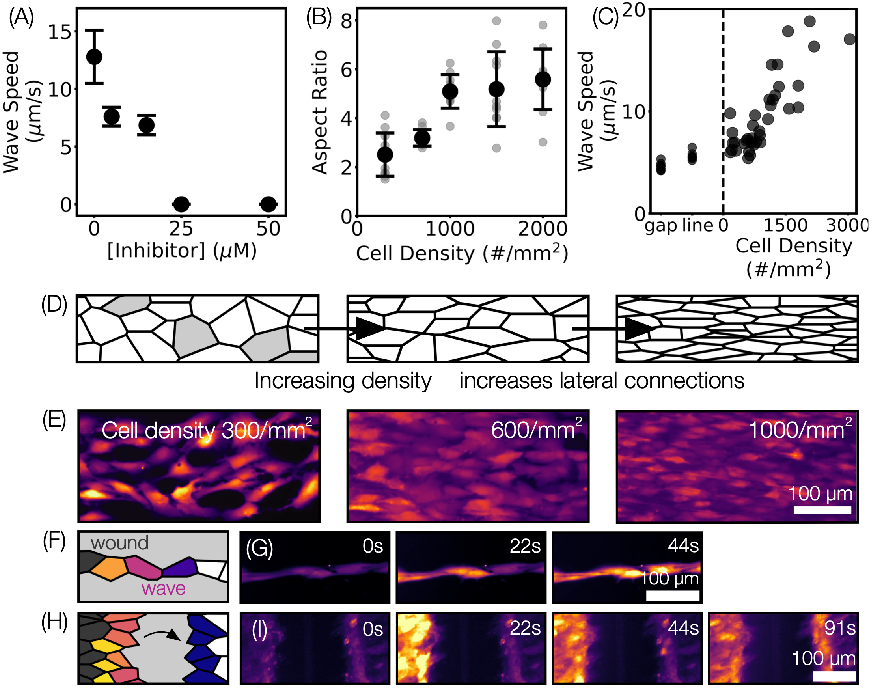
Gap junction inhibitors and cell density modify intercellular coupling strength and wave speed. (A) Tissues treated with varying concentrations of Meclofenamic Acid (MA), a gap junction inhibitor. Wave speed slowed and then was abrogated with increasing concentrations of MA in scratches oriented in the *θ* = 0 direction. (B) Aspect ratio of cells increased with increasing density. (C) Velocity of the calcium wave in scratches oriented at 0 increased with cell density. (D) Schematic representing that increasing density in tissues increases lateral contacts. Grey squares represent areas with no cells. Lower density tissues are not confluent. (E) RPE-1 cells at 3 different densities. As the cells become denser there is less free space between cells and more cell-cell contact. Schematic (F) and times series (G) of wave propagation across a 1D line of cells in the 0 direction. Schematic (H) and time series (I) of wave propagation across a gap of no cells in the 0 direction.

Upon measuring that decreased gap junction activity slows the wave, we asked if conversely increasing cell-cell connectivity would trigger increased wave speeds. We increased the amount of cell-cell connectivity by modulating tissue density. From a purely geometrical argument, increasing the density of packed particles increases the average number of contact points between neighbors. Increasing the anisotropy of packed particles preferentially increases contact points laterally between neighbors. We measured that as tissues get denser, cell anisotropy also increased, highlighting that cells were getting crowded together laterally (Fig. 3*B,D*).

We performed linear wounds oriented at 0 radians in tissues of varying density. The calcium wave moved faster in denser tissues (Fig. 3*C*). This revealed that a global wave moving along a cell’s long axis is sped up by increased lateral connections along a cell’s short axis.

To test the limits of wave propagation through cell-cell contact we performed linear wounds on single-file lines of RPE-1 cells (Fig. 3*G*). In this condition, only a minimal contact existed between cells at their tips, compared to our experiments in dense tissue that had lateral and tip contacts with neighbors. The wave moved slower in the line configuration than in the 2D tissues (Fig. 3*C*) but it was still able to propagate.

Finally, to further confirm the impact of cell-cell coupling on wave speed, we performed a linear wound experiment on a tissue that had a gap between two populations of aligned cells. Consistent with published results, [1] we saw that a wave triggered in one population could stimulate a calcium wave in the second population (Fig. 3*I*). However, the time it took for the unconnected population of cells to respond and light up was slower than the time for the signal to propagate in contacting cells (Fig. 3*C*). This demonstrated that while gap junctions mediate a fast signal between cells, a diffusible signal can also stimulate a wave moving over longer timescales. We did not observe an extracellular diffusion-driven wave when inhibiting gap junctions, suggesting a need for some cell-cell junction activity to respond to the extracellular signal.

We turned to modeling to develop a more quantitative understanding of how calcium waves move in our structured tissue. We modify a well-studied model of nonlinear front propagation, the KPP equation [24]

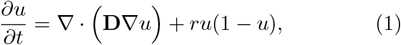

where **D** is the diffusion tensor, *r* is the reaction rate, and *u* is a scalar field, which we take to represent the calcium concentration. Upon perturbation to a uniform state *u* = 0, the system rapidly transitions to a stable value *u* = 1. Spatially-localized perturbations propagate outward as a well-defined solution with a characteristic velocity 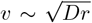 [25]. In this simplified framing of the experimental dynamics, intercellular coupling mediated by gap junctions and extracellular communication through the media are collectively expressed by the anisotropic diffusional transport of *u*, shown schematically in (Fig. 2*A*), while dynamics leading to the rapid spike in intracellular calcium are approximated by the nonlinear reaction *ru*(1*− u*). We express the diffusion tensor as follows

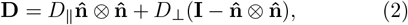

where *D*_*I*_ and *D*_*⊥*_ are the diffusivities parallel and perpendicular to the nematic director 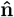, respectively, and *⊗* represent outer products of the units vector. The nematic director 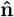 is a headless vector encoding the orientation field of the tissue. Parallel and perpendicular diffusivities have successfully described anomalous Brownian motion in nematic liquid crystals [26–29], however, extending a nematic-like diffusion tensor to a reactiondiffusion system is unexplored in the context of epithelia to our knowledge.

Before presenting predictions of wave propagation by numerically integrating Eqn. 1, we first estimate the impact of anisotropy on wavespeed analytically. In the sharp interface limit, neglecting the chemical wavefront’s curvature, and assuming uniformity in the underlying nematic, the wavefront velocity is given by

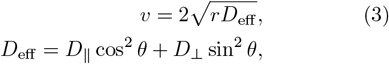

where the effective diffusivity depends on the angle 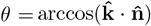 between the wavevector 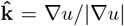 and the nematic director 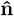. We show the full derivattion of Eqn. 3 in the supplement Eqns. S1-S10. We fit Eqn. 3 to the experimental data in (Fig. 1*I*) and find that it captures the leading-order effect: maximal velocity perpendicular to the nematic director and minimal velocity parallel to the nematic director. Using two free fit parameters for *rD*_∥_ and *rD*_⊥_, we obtain the ratio of the diffusivities *D*_⊥_/*D*_∥_ *≈* 4.2. Because our fit finds that the perpendicular diffusivity is roughly 4 times the parallel diffusivity, the perpendicular wave speed *v*(*θ* = *π*/2) should be roughly twice the parallel wave speed *v*(*θ* = 0), which is exactly what we observe in experiment (Fig. 1*I*). We attribute deviations from this idealized theory in (Fig. 1*I*) to the complex microstructure of the tissue, which we discuss further in the conclusion.

Our purely nematic tissues seeded on grooved substrates allowed us to identify the relationship between a cell’s orientation with respect to a calcium wave and the global wave speed. However, nematic defects exist in the tissue where regions of cells are locally aligned but globally disordered. In unconfined RPE-1 tissue nematic defects were separated by 500 - 1000*μ*m with the distance between defects decreasing with increased density consistent with published studies [22]. Important biological processes are known to localize at nematic defects, such as apoptosis [8] and budding [20, 30]. These events can also trigger calcium signaling [13–15].

We therefore leveraged our model to simulate the evolution of a calcium wavefront from a wound near a topological defect. We integrate Eqns. 1, 2 subject to circular and linear initial conditions to emulate experimental linear and point wounds. Simulations of a linear wound in a uniformly aligned nematic (Fig. 4*A*) recover similar linear translation of the chemical wavefront as was observed in experiments (Fig. 1*F*,*H*). Similarly, a point wound in a uniform nematic (Fig. 4*D*) assumes an elliptical-like wavefront shape, as was found in experiment (Fig. 1*K*). However, introducing topological defects dramatically changes the behavior of chemical waves. An initially flat plane wave traveling through a topological defect does not remain flat. As the wave travels, the nematic director around it changes — hence changing the angle between the wave vector and nematic — altering the local wave speed and distorting the wavefront (Fig. 4*B*,*C*).

**FIG. 4.**
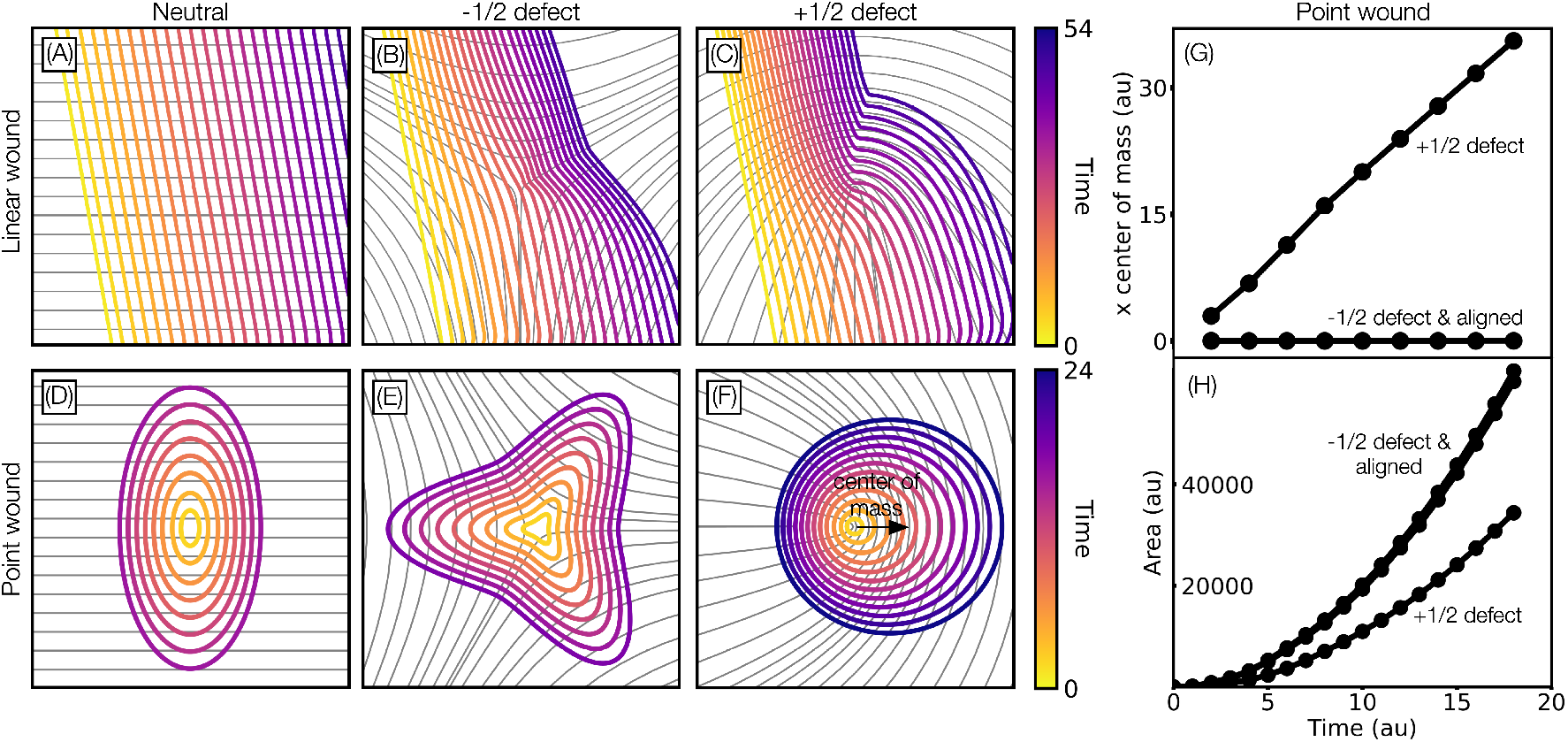
Nematic texture gives shape to chemical wavefronts. We simulate the evolution of a chemical wave traveling through different nematic textures with the experimentally measured ratio of perpendicular to parallel diffusivities *D*_⊥_*/D*_∥_ ≈ 4. (A) A plane wave maintains its shape and travels at a constant speed through a uniformly-aligned nematic. (B,C) However, a plane wave traveling through a (B) − 1/2 and (C) +1/2 topological defect has its wavefront distorted. The angle between the wavevector and nematic director changes while the wave travels, hence modulating the local wavespeed. (D) A point source emitted outward in a uniformly-aligned nematic assumes an elliptical wavefront shape. (E,F) Wavefronts from point sources in topological defects encode the symmetry of the defect. (E) A wavefront emitted outward from a − 1/2 defect has trefoil symmetry. (F) And a wavefront emitted outward from a +1/2 defect assumes a circular shape and translates to the right. (G) For the point source emitted outward in a +1/2 defect, the *x*-coordinate center of mass increases linearly whereas the point sources emitted from a − 1/2 defect and a uniformly aligned nematic do not translate. The *y*-coordinate center of masses do not translate. (H) The area enclosed by the wavefront increases faster from a point source emitted in a − 1/2 defect and uniformly aligned nematic than from a point source emitted from a +1/2 defect.

We observed a symmetry-preserving relationship between nematic defects and the chemical wave front’s shape when initiated from a point wound coinciding with the defects’ cores. We find that a wavefront emanating from a − 1/2 defect exhibits a trefoil shape following the threefold symmetry of the underlying nematic (Fig. 4*E*). And moreover, a wavefront emanating from a +1/2 defect displays a nearly circular shape with the centroid acquiring a steady shift of the contour’s center of mass to the right as the region emanates outward (Fig. 4*F*,*G*). The broken polar symmetry of +1/2 defects is well-studied in active nematohydrodynamics, but their impact on front propagation has not been documented to our knowledge. We further quantify the areas of the growing fronts and find the area enclosed by the wavefront grows at different rates depending on the tissues underlying nematic orientation (Fig. 4*H*). The contour area grows slowest for a point source emitted from a +1/2 defect compared to a point source emitted at a − 1/2 defect or in an aligned tissue.

## DISCUSSION

Biology self-organizes by closely coupling form, mechanics, and biochemistry. Deepening our understanding of how these physical processes interrelate across scales and cell types is an ongoing effort that will impact developmental biology and medicine. Here, we experimentally demonstrate that an interplay exists between the speed of calcium wave propagation and the tissue’s microstructure, specifically its nematic order. We conclude that the origin of wave speed anisotropy is the local coupling environment of cells, wherein elongated cells in a tissue more strongly couple to their lateral neighbors due to increased contact area. We developed a simple mathematical model that recapitulates our observations of how wave speed depends on this structure. While our study focuses on the propagation of calcium waves through epithelial media, the underlying physical mechanism is generic and likely to apply broadly to other cell types and communication pathways. The impact of anisotropy could also be significant even in the absence of coherent, large-scale signaling waves, as we’ve studied here.

The analogy between tissues and nematic liquid crystals has provided insight into collective dynamics of cells. Liquid crystals are birefringent whereby the material has two different refractive indices for polarized light either parallel or perpendicular to the material’s internal molecular orientation. We demonstrate birefringence in epithelial tissues with a calcium wave moving at different speeds depending on its orientation relative to a tissue’s nematic director. Our results strengthen the analogy of tissues as nematic liquid crystals.

We demonstrate that nematic defects desynchronize a propagating calcium wave. The area inside the calcium wave contour grew at different rates, depending on the underlying tissue’s nematic orientation. Therefore, cells at the same spatial distance from the point wound received information at different times depending on if they were triggered on aligned tissues, +1/2 or −1/2 defects. The same mechanical wounding leads to different tissue-scale outcomes. Similarly, in propagating plane waves, the tissue structure distorts the wavefront, and cells equidistant from the wave’s initial starting point experience a wave passing at different times depending on their position relative to a nematic defect. This asynchrony in when neighboring cells receive a calcium signal could have functional importance *in vivo*. A calcium signal can trigger cells to constrict [31], and temporal variation in cell constrictions is important in guiding diverse shape changes during development [32].

By introducing nematic order-dependent diffusivity, we predict the dependence of wave speed on the relative angle between the wave front and director. While our model qualitatively shows that maximal and minimal wave speeds, respectively, correspond to waves moving perpendicular and parallel to the director, Fig. 1*I* shows quantitative differences between theory, Eqn. 3, and observations at intermediate angles. We posit that this discrepancy is due to the complexity of the tissue’s microstructure. For example, while the cells are elongated and aligned with their neighbors, the disordered packing of the tissue exhibits a highly hexatic structure in addition to nematic order (Fig. S1*J*). Recent studies of epithelial flows have quantified cellular shape and packing order parameters, incorporating them into active hydrodynamic theories [33–35]. Our work suggests that this structure also influences cellular communication. Wave speed anisotropy might even serve as a measure of the underlying structure. Taken together, our observations suggest an exciting line of experimental and theoretical inquiry that further investigates the relationship between wave propagation and the various forms of order that exist in tissues.

In our experiments, we observed an overall decrease in the speed of point-wound-initiated waves compared to scratch wounds. This finding was consistent with the literature, which noted that the amount of initial wounding impacts wave speed [36]. We hypothesize that the decreased contrast in wave speed between parallel and perpendicular directions could be related to the diffusible signal that is released from the wound. Previous work has shown that calcium wave dynamics depend on the availability of extracellular ATP, which plays an excitatory role in the signaling dynamics. ATP is released endogenously by cells during calcium signaling but can also be introduced exogenously as an experimental control parameter [1, 37]. In some cell types, the absence of a baseline level of extracellular ATP prevents the propagation of signals altogether [1]. Cell injury releases freely diffusing ATP and contributes to the initial excitation. The geometry of the wound significantly influences this initial distribution. Comparing linear and point wounds, an initial circular distribution of ATP will diffuse radially and dilute more rapidly than a linear distribution. Thus, we posit that the slower wave speeds we observe are due to this extracellular effect. This is further corroborated by a wave speed dependence on initial wound size, with larger wounds.

Capturing these physical processes theoretically is beyond the capabilities of our simplified model, which evolves a single scalar value over time and omits biochemical details in favor of a focus on anisotropic diffusion. A future modeling direction is to explicitly capture the transport of specific molecular species, such as Ca^2+^, ATP, and IP_3_ [38–40]. In such a model, one could mimic experimental techniques by shaping initial extracellular ATP distributions.

Examining the coupling between topology and chemical processes is an active area of focus with researchers investigating both how chemical perturbations can precipitate structural rearrangements and how topology can shape reaction-diffusion processes [10, 41, 42]. In particular, nematic defects localize important biological processes in various tissues, including cell extrusion in biofilms [8, 20, 30] and limb growth in multicellular life [10, 41, 43]. *In vivo* calcium waves are commonly observed emanating from such events [13]. Beyond animal epithelia, plant tissue also exhibits chemical-topological feedback loops. For example, the growth hormone auxin is pumped in a directional manner along the axis of cells, which, in turn, grow to orient along those dynamic gradients [44–46]. This feedback loop reinforces the gradients and spontaneously colocalizes auxin maxima with topological defects. Another example of chemical feedback in polar media is motile Dictyostelium cells, which use cyclic AMP signaling to collectively select a gathering site and cooperatively form multicellular structures [47]. Taken together, these examples highlight how structural–chemical feedback mechanisms organize biological pattern formation. Our work suggests that anisotropic diffusion is another tool used by biological systems to shape collective organization.

## CONCLUSION

Our work uncovers an interplay between cell nematic orientation and calcium wave speed. We show how these two tissue-scale cues interact to impact long range communication in epithelial tissues. Our work therefore provides an entirely new perspective on how physical and biochemical cues can be integrated to coordinate collective cell behaviors. This mechanism has relevance for the diverse range of tissues that display calcium waves during development and disease.

## AUTHOR CONTRIBUTIONS

H.G.Y. and A.C.W. designed research; A.C.W., B.C.S, and M.M.N. performed research; all authors analyzed data; all authors wrote the paper;

## ACKNOWLEDGMENTS

Research reported in this publication was supported by the National Institute Of General Medical Sciences of the National Institutes of Health under Award Number R00GM136915. The content is solely the responsibility of the authors and does not necessarily represent the official views of the National Institutes of Health. We thank Dr. Piali Sengupta for the RPE-1 cells. The development of the computational model was supported by Department of Energy BES No. DE-SC0022280 (MMN). We also acknowledge the use of the optical and biomaterial facilities supported by NSF MRSEC Grant No. DMR2011846, the Brandeis Light Microscopy Core Facility RRID: SCR 025892 for equipment and Dr. Andrew Stone for assistance.

## MATERIALS AND METHODS

### Cell culture

RPE-1 Cells were maintained in Dulbecco’s Modified Eagle Medium (DMEM) with 1000 mg/L glucose, L-glutamine, and pyruvate (Thermo Fisher, cat#: 11885092), supplemented with 10% (v/v) fetal bovine serum (Fisher Sci., cat#: MT35011CV) and 100 μg/mL penicillin-streptomycin (Sigma, cat#: P4333). Cultures were incubated at 37 °C in a humidified atmosphere with 5% CO_2_. All experiments were performed using cells at or below passage 30.

### Substrate preparation

#### Fabricating Nematically Aligned Tissues

Microgrooves were created on the base of surface-treated, sterile 6-well tissue culture plates (Sigma, cat#: CLS3516) by gently dragging 2000-grit sandpaper across the well surface using light thumb pressure. Cells were seeded onto the grooved substrates and allowed to grow until confluence over several days. As cells adhered and proliferated, they aligned along the direction of the grooves, forming a tissue with local long-axis alignment. To vary cell density, the culture duration was varied: shorter growth periods produced lower densities, while extended growth periods yielded higher densities. For imaging cells with confocal microscopy (Fig. 2*C,D*; Fig. S2*A,C*) cells were seeded into 35 mm uncoated glass bottom dishes (Fisher Sci., cat#: NC1844456) and allowed to grow to confluence. For immunofluorescence staining, cells were seeded onto a cover slip and allowed to grow to confluence.

#### Preparing a 1D Line of Cells

To generate 1-D lines of cells, cells were seeded at very low density onto the microgrooved substrates described above and incubated for 24–48 hours. After this period, wells were screened to identify those in which cells had aligned and proliferated along a single microgroove, forming a continuous line of cells. Approximately 10% of wells exhibited this pattern and were selected for further experiments. Wounds were scratched at the edge of one of these lines.

#### Preparing a Gap Within a Tissue

To create a gap within cell tissues, a silk sewing pin (Amazon) was used to create a wound across the aligned tissue and then washed to remove dead cells. The plates were then labeled with Calbryte-520AM and subsequently used in experiments where a scratch was created within 500 μm of the gap.

### Reagents

#### Intracellular Calcium Labeling

Intracellular calcium was visualized using Calbryte-520AM (Fisher Sci. Sci., cat#: NC1812243). The dye was initially dissolved in DMSO and subsequently diluted to a final concentration of 5 μM in Hanks’ Balanced Salt Solution supplemented with calcium and magnesium (HBSS + Ca; Fisher Sci., cat#: J67763.). Probenecid (Sigma, cat#: P8761) was also added to this solution for a final concentration of 1mM. Cells were incubated with 1 mL of the 5 μM Calbryte-520AM solution per well for 1 hour at 37 °C. Following incubation, cells were washed with phosphate-buffered saline (PBS) and then maintained in HBSS + Ca. Imaging was performed immediately after dyeing, with HBSS + Ca used as the imaging medium unless otherwise specified.

#### Gap Junction Inhibition

Meclofenamic acid (MA; Thermo Fisher, cat#: J60484) was first dissolved in sterile deionized (DI) water to prepare a 1 mM stock solution. This stock was further diluted in HHBS + Ca to yield working concentrations of 5 μM, 15 μM, 25 μM, 35 μM, and 50 μM, as required for the experiments. Dishes were first incubated with Calbryte-520AM for 1hr, then washed before being incubated with MA for 20min prior to imaging without washing.

#### Immunofluorescent Staining

Cover slips containing confluent tissue cultures were fixed with 4% paraformaldehyde (Fisher Sci., cat# AAJ19943K2) for 15 minutes. Between each step, cover slips were washed 3*×* in PBS. Following fixation, cells were permeabilized using 0.1% Triton X-100 (Fisher Sci., cat#: PI28314) for 10 minutes. Non-specific binding was blocked using 1% bovine serum albumin (BSA; Fisher Sci., cat#: AAJ61089AP) in PBS for 30 minutes.

Samples were then incubated with a 1:1000 dilution of Anti-Connexin 43 Antibody produced in rabbit (Sigma, cat#: C6219) overnight at 4°C in the dark, using a box lined with wet paper towels. The following day, a 1:500 dilution of Anti-Rabbit IgG (H+L), CF™568 antibody produced in chicken (Sigma, cat#: SAB4600426) and a 1:400 dilution of Phalloidin (Thermo Fisher, cat#: A12379) were applied for 1 hour in the dark. Cover slips were mounted onto glass microscope slides using Pro-Long™ Diamond Antifade Mountant with DAPI (Thermo Fisher, cat#: P36966) and left to cure in the dark for 24 hr at room temperature. Imaging was performed using a confocal microscope, as described in the Microscopy and Imaging section.

### Tissue wounding

The cells were wounded during image acquisition on the microscope. A silk sewing pin (Amazon) was glided along the cell surface to create linear wounds. Point wounds were made by lightly touching the cell surface with the ball end of the pin.

### Microscopy and Imaging

For all experiments, wounds were created during image acquisition and acquisition continued for at least several minutes up to 10 minutes.

Fluorescence and phase contrast imaging was performed using a Nikon Ti2 inverted microscope equipped with Nikon Elements software for image acquisition and an OKO CO_2_ environmental enclosure to maintain physiological conditions. Fluorescence excitation for wounding experiments was achieved using a 480 nm excitation line via the GFP setting of the Sola Light Engine. Immunofluorescence excitation was achieved using a 480 nm excitation line via the GFP setting (actin), a 560 nm excitation line via the mCherry setting (Connexin-43), and the DAPI setting (Nuclei). Images were acquired using a Nikon Plan Fluor 4 *×* /0.13 NA objective (WD 17.2 mm), a Nikon Plan Fluor 10 *×* /0.30 NA Ph1 DLL objective, or a Nikon Plan Fluor 20 *×* /0.5 NA 0FN25 Ph1 DLL objective. Phase contrast imaging was performed using the 10*×* objective to visualize cell morphology. Immunofluorescent images were captured with the 20*×* objective. Time-lapse imaging was conducted at a frame rate of 500 ms per frame. Images were captured using a Hamamatsu ORCA-Flash4.0 LT digital CMOS camera (model C11440).

Confocal imaging was performed using a Nikon AX-R system mounted on a Ti2-E inverted microscope (Nikon Instruments). Samples were maintained under environmental control during imaging at 37°C and 5% CO_2_ using a stage-top incubator. Excitation was achieved using a 488 nm diode laser at 25% laser power. A Nikon 40 *×* /1.15 NA water-immersion objective was used with deionized water as the immersion medium. The resonant scanner was used for image acquisition. The pixel size was 0.12 *μ*m. The pinhole was set to 2.1 Airy units. Emission was collected over a spectral detection window of 499.0 - 551.0 nm using a GaAsP detector with a gain setting of 1.5. Time-lapse imaging was conducted at a frame rate of 250 ms per frame.

### Image Analysis

Orientation fields were analyzed using the OrientationJ plugin for Fiji, which calculates local orientation and coherence based on the structure tensor method [48, 49]. A custom python script was used to overlay the orientations onto the phase contrast images to display the distribution of the orientation angles.

To measure wave propagation speed in response to linear wounds, kymographs of signal intensity were generated in Fiji (Fig. S1*B*). To generate kymographs, a rectangular region of interest (ROI) was drawn along the axis of interest using Fiji (Fig. S1*A*). The “Reslice” tool was applied from top to bottom, generating a timelapse stack of the intensity along the ROI. A Z-projection (average intensity) of the resliced stack was then computed to produce the final kymograph (Fig. S1*B*). An intensity threshold was applied to the kymographs to enhance contrast between background and wavefront signal (Fig. S1*C*). The leading edge of the kymograph was then fit to a straight line. The inverse of the slope of this line was taken as the wavefront speed (Fig. S1*D*).

For point wounds, a separate custom Python script was used to track wavefront progression over time. Two selected frames were binarized (Fig. S1*F*,*G*), and the earlier frame was subtracted from the later one to generate a ring representing the spatial extent of the wavefront displacement during that time interval. Small objects were removed to eliminate noise, and a light dilation operation was applied to fill small holes and smooth the wavefront (Fig. S1*H*). Radial widths of the resulting ring were then measured as a function of angle. Wave speed at each radial angle was computed by dividing the radial width by the time interval between the two frames (Fig. S1*I*).

Intensity profiles were measured using a custom Python script by drawing a horizontal line across the center of the cell of interest. Fluorescence intensity along this line was tracked over time to generate intensityversus-time data. In some cases, the intensity values along the line were averaged at each time point to produce Intensity vs. Time plots (Fig. 2*E*,*F*). In other cases, the full intensity profile along the line was retained to generate Distance vs. Intensity plots (Fig. S2*B*,*D*).

Aspect ratios of individual cells were determined in Fiji by measuring the lengths of the long and short axes. Ten cells were randomly selected per image for these measurements. Cell density was quantified by drawing a small rectangular region of interest (ROI) on the image using Fiji. The number of cells within this ROI was manually counted and divided by the ROI area to obtain density values. To analyze local tissue organization in (Fig. S1*J*) cell centroids were manually marked in each field of view. These centroid coordinates were input into a custom Python script to generate Voronoi tessellations, from which neighbor relationships were extracted.

### Statistical Analysis

Statistical analyses were performed using Python. A paired *t* -test was used to compare parallel and perpendicular speeds of point wounds. Statistical significance was considered at *p <* 0.001. All experiments were performed with at least three replicates, measuring wave fronts propagating over a minimum distance of 250 μm from the wound. Where applicable, error bars represent the standard deviation.

### Simulation Details

We simulate Eqns. 1 using the finite element solver COMSOL. We arbitrarily set *r* = 1 and integrate the equations on a 400*×*400-sized square domain for 30 time units. We set *D*_∥_ = 0.5 and *D*_⊥_ = 2, which corresponds to a velocity *v*_⊥_/*v*_∥_ = 2 and mirrors the experiment, Fig. 1*I*. When simulating the dynamics of point-wound and scratch-wound-initiated waves interacting with defects, we avoid singularities in 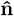 by smoothly reducing the anisotropic contributions of the diffusion tensor to zero via a Gaussian divot with unit standard deviation located at the defect core. This region is small compared to the domain size. Thus, the large-scale dynamics we display are largely unaffected by this regularization.

## Supporting Information Text

## Model

### Minimum wavespeed

Here, we derive the expression for the minimum wavespeed predicted by the anisotropic KPP equation, which we repeat here for clarity

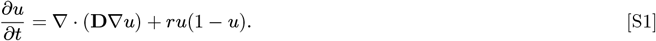

The variable *u* is the chemical concentration, **D** is the diffusion tensor, and *ru*(1 *− u*) is the nonlinear reaction term responsible for the model’s excitability, which possesses unstable and stable fixed points at, respectively, *u* = 0 and *u* = 1. To find an expression for wave speed in anisotropic but spatially uniform media and negligible wavefront curvature, we consider a solution of the form

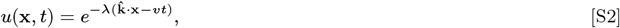

where *v* is the wavespeed of a plane wave, *λ* is an unspecified spatial decay length such that when λ *>* 0 the wave decays ahead of the propagating region, and 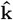 is the wavevector in the direction normal to the chemical field 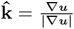. Retaining only terms linear in *u* and substituting the form Eq. (S2) into Eq. (S1) gives

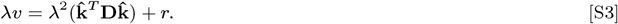

Solving for *λ* gives

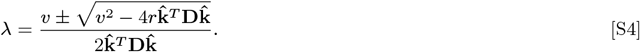

For our initial assumption that λ ∈ ℜ to hold, we find the condition that

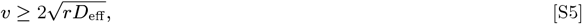

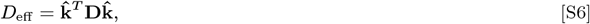

where *D*_eff_ is the effective diffusivity in the direction of propagation 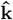. In the case of isotropic diffusivity, **D** = *D****I*** and we recover the classic result

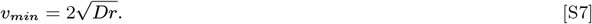

### Nematic diffusivity

Experiments observe chemical signals travel faster perpendicular than parallel to the nematic director. To model this behavior, we assume the diffusion tensor has two components: one parallel to the nematic director ***n*** and another perpendicular to the nematic director

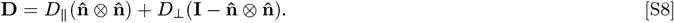

where *D*_∥_ is the diffusivity parallel to the nematic director and *D*_⊥_ is the diffusivity perpendicular to the nematic director. The effective diffusivity in the direction of the wavevector ***k*** is

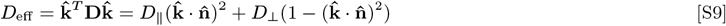

Let 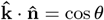. From Eq. (S5) and Eq. (S9), we find the minimum wavespeed in a nematic

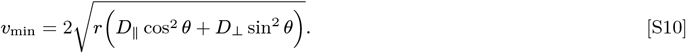

**Fig. S1.**
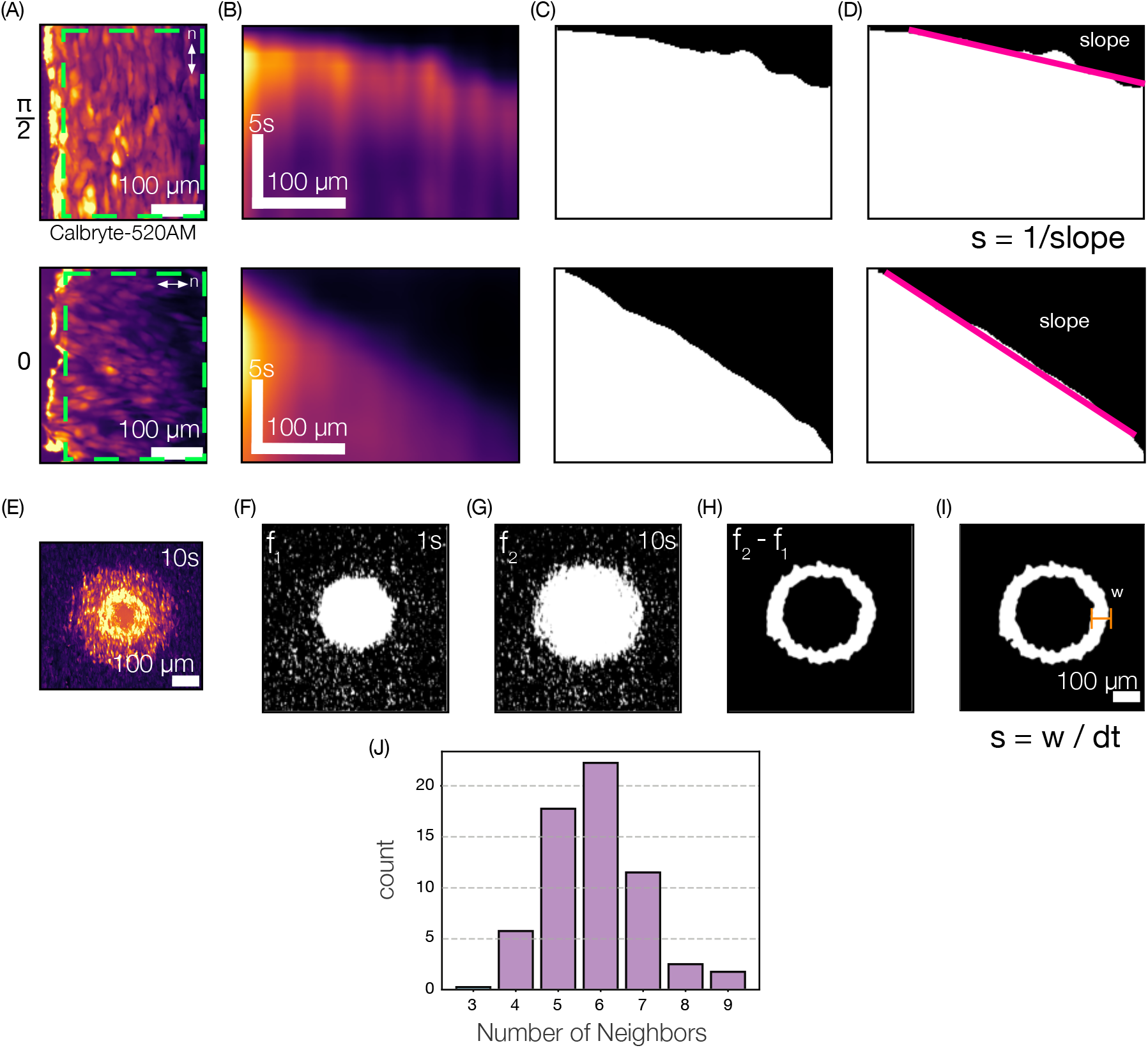
Measuring calcium wave speeds. (A-D) Measuring Speeds for Scratch Wounds. (A) Fluorescence image of a linear wound after scratch, 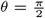 (top) and *θ* =0 (bottom). Green rectangle represents the area that was resliced to created the kymograph. (B) Kymographs resulting from reslicing (A) from top to bottom and taking the average intensity z-projection, showing wave propagation for 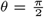 (top) and *θ* =0 (bottom) cases. (C) Kymograph after thresholding out noise in Fiji to remove background, for 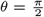 (top) and *θ* =0 (bottom). (D) Speeds were measured by taking the slope of the threshold image and taking the inverse. (E-I) Measuring Speeds for point wounds. (E) Fluorescence image for point wound. (F) Binarized frame 1 for speed measurement of point wound. (G) Binarized frame 2 for speed measurement of point wound. (H) Subtraction of frame 1 from frame 2 gives the change of the wave front between frame 1 and frame 2. (I) Measuring how the radial width of this wave front change allows speed measurements for every polar angle, *ϕ*, to be calculated. (J) Distribution of number of cell neighbors in an aligned RPE1 tissue.

**Fig. S2.**
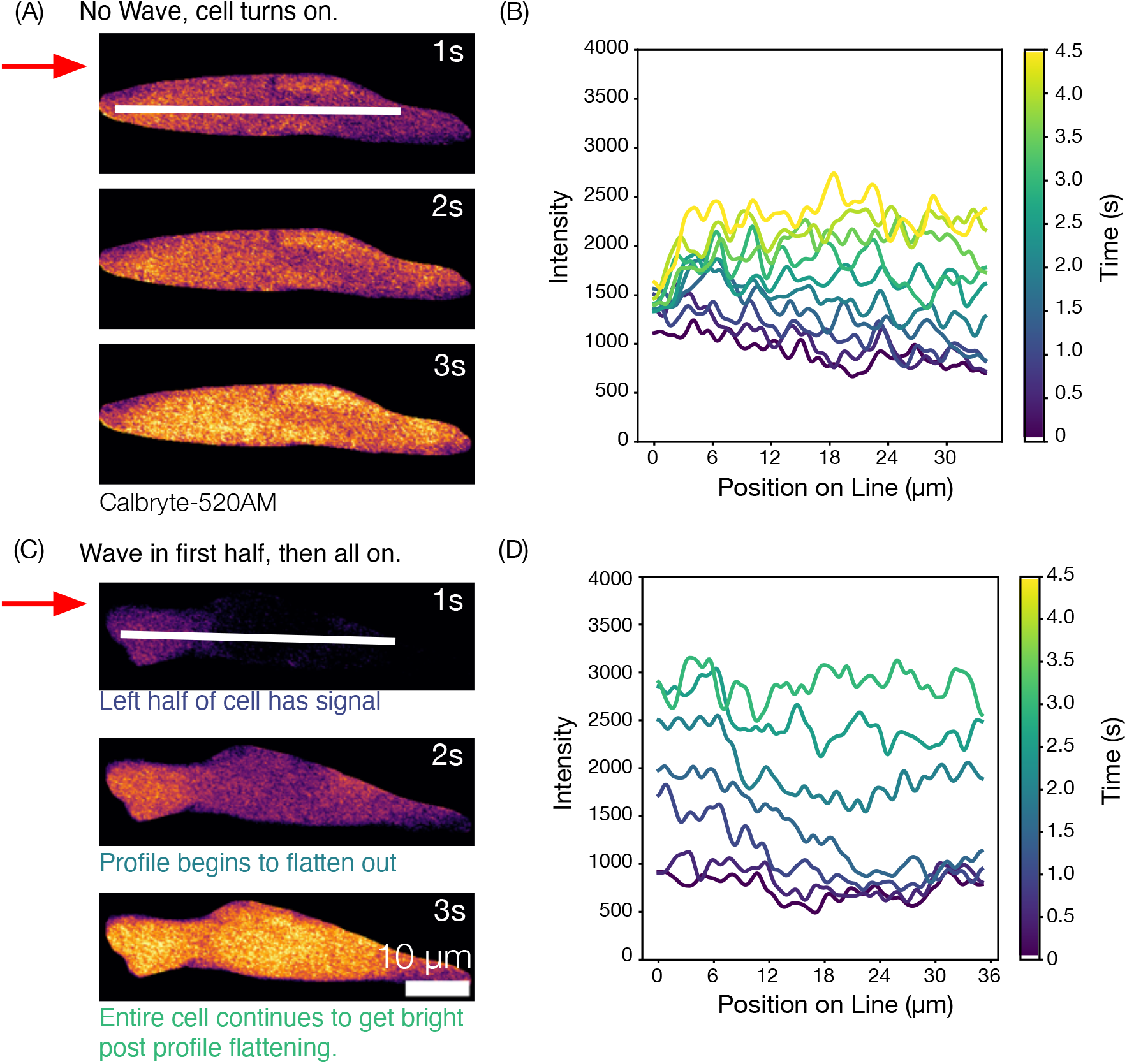
Cells either turn on all at once or show a wave in one region of cell before brightening across the whole cell at once. (A) Time series of cell turning on all at once. White line indicates where the intensity profiles were measured at. Red arrow shows global wave direction. (B) Intensity profiles over time measured at the white line in (A). Profiles show that over time the cell is brightening along the entire line, representing the cell turning on rather than a wave. (C) Time series of cell showing a wave until the halfway point, then turning on at once. (D) Intensity profiles over time measured at the white line in (C). At t= 1.0s, the intensity profile shows higher intensity in the left side of the cell. By t=2.0s, the profile has begun to flatten out. By t= 3.0s, the profile had flattened out and the entire cell’s brightness had increased. Montage images were taken from tissues and were masked to only show the cell of focus.

**Fig. S3.**
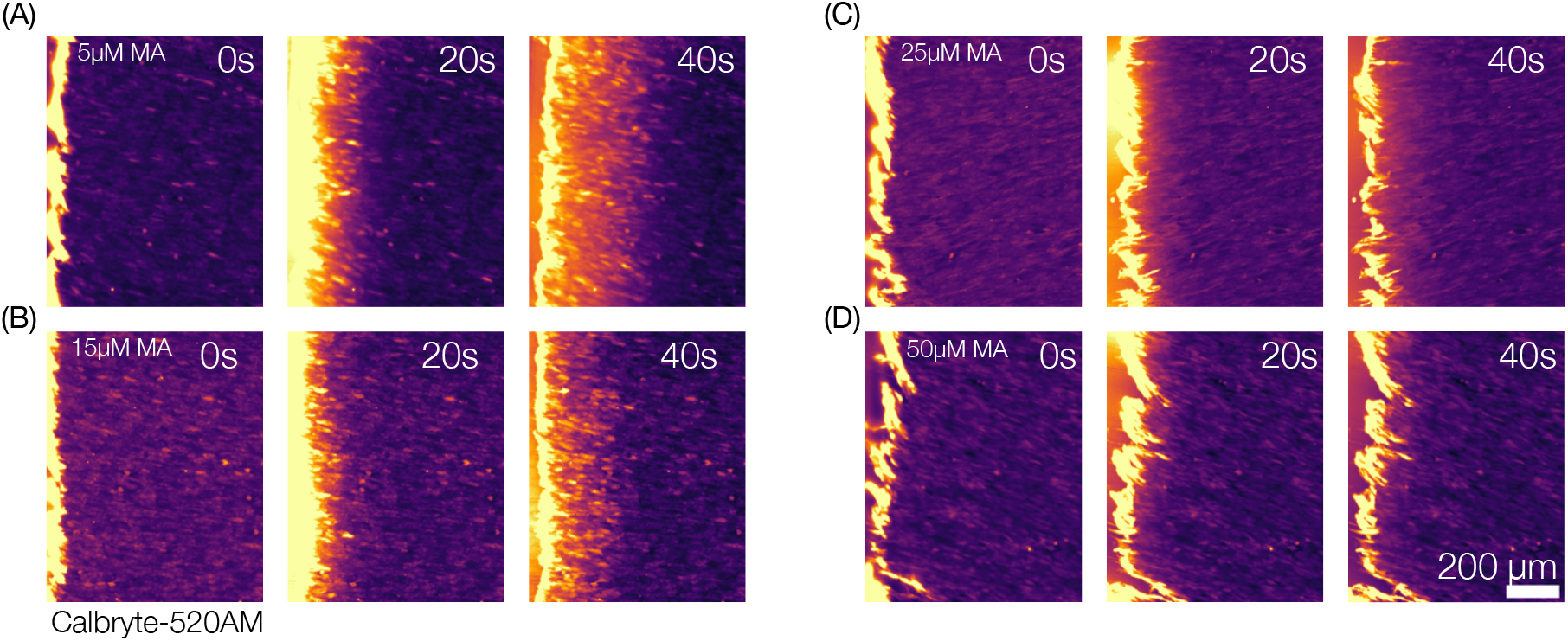
Meclofenamic Acid (MA) a Connexin-43 Inhibitor blocks propagation of calcium waves at high dosage. (A-B) 5*μ*M and 15*μ*M MA, respectively, still show calcium wave propagation, however the 15*μ*M case has a slower wave than the 5*μ*M case. (C-D) Concentrations at and above 25*μ*M halt wave propagation, showing only wounded cells expressing Ca2+.

**Movie S1. Time lapse of calcium wave propagating across the cell’s short axis**, 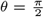 **(A) and long axis**, *θ* = 0 **(B) resulting from linear scratch wound**.

**Movie S2. Time lapse of calcium wave propagating radially away from a point wound**.

**Movie S3. Time lapse of calcium wave propagating across two lateral neighbors. Cell outlined in green signals first, and cell outlined in blue signals second**.

**Movie S4. Time lapse of calcium wave propagating across two tip to tip neighbors. Cell outlined in green signals first, and cell outlined in blue signals second**.

**Movie S5. Time lapse of calcium wave propagating across a (A) 1D line of cells and (B) across a gap of no cells. The wave travels faster along the 1D line than across the gap. The line video stops after 44s, while the gap continues through 91s**.

**Movie S6. Simulation of a plane wave traveling through a uniformly aligned nematic, a** −1/2 **topological defect, and a** +1/2 **topological defect**.

**Movie S7. Simulation of a point source emanating outward in a uniformly aligned nematic, a** − 1/2 **topological defect, and a** +1/2 **topological defect**.

